# In search for the invisible: motor inhibition in monkey premotor cortex and its RNN replicas

**DOI:** 10.1101/2025.11.24.690225

**Authors:** Nicola Alboré, Andrea Galluzzi, Pierpaolo Pani, Stefano Ferraina, Maurizio Mattia

**Affiliations:** Natl. Center for Radioprotection and Computational Physics, Istituto Superiore di Sanitá, Rome, 00161, Italy; Research Center “Enrico Fermi”, 00184 Rome, Italy; PhD program in Physics, “Tor Vergata” University of Rome, 00133 Rome, Italy; Dept. of Physiology and Pharmacology, “Sapienza” University, 00185 Rome, Italy

## Abstract

Controlling actions in dynamically changing environments requires flexible and efficient motor control. A fundamental challenge in neuroscience is to uncover how cortical circuits generate, adjust, and sometimes suppress planned movements. To address this, we combined recordings from dorsal premotor cortex (PMd) of macaque monkeys performing a stop-signal task, with a recurrent neural network (RNN) model inferred directly from multi-unit activity. This data-driven “digital twin” reproduced the cortical population dynamics underlying motor planning and inhibition, revealing how internal network states shape behavior, and generating synthetic neural trajectories for unseen conditions. RNN internal state explained reaction time fluctuations across trials, reflecting stochastic components of motor readiness and endogenous variability of PMd activity. The same pre-Go latent state also constrained movement inhibition modulating the network’s response to Stop signals by reshaping the attractor dynamics of a null-potent subspace. These results establish a mechanistic link between latent cortical dynamics and flexible behavioral control, demonstrating how autonomous RNN inference can uncover circuit-level computations.

## Introduction

The dorsal premotor cortex (PMd) occupies a fundamental role in the cortical motor hierarchy, integrating sensory information, contextual cues, and internal goals to shape voluntary movement planning and execution [1, 2, 3]. Recent studies have emphasized that PMd activity does not simply reflect discrete stimulus-response mappings but unfolds dynamically across low-dimensional neural manifolds corresponding to holding, planning, and execution stages of motor behavior [4, 5, 6, 7, 8]. In stop-signal tasks, where subjects are instructed to prepare and sometimes cancel planned movements based on an unpredictable “Stop” signal [9, 10], PMd trajectories exhibit early divergence between no-stop and correct-stop trials, suggesting that motor suppression is implemented via a redirection of ongoing neural trajectories rather than through a late-stage inhibitory override [11, 12, 13, 6].

Single-trial analyses have shown that the probability of successful inhibition depends strongly on the internal neural state at the time of the Go cue, with specific subspaces within PMd associated with different behavioral outcomes [14, 6]. These findings propose a view of PMd as a dynamical system, wherein internal geometry constrains the potential evolution of neural trajectories and ultimately shapes behavioral responses. However, despite increasingly rich datasets capturing high-dimensional population activity, current analytical approaches remain largely observational. Dimensionality reduction techniques such as principal component analysis (PCA) or demixed PCA (dPCA) reveal geometric structure but lack the ability to simulate how trajectories autonomously evolve over time [15, 16]. Similarly, decoding approaches can predict behavior from neural data but do not model the causal unfolding of internal cortical dynamics. Thus, there remains a pressing need for frameworks that move beyond static or feedforward descriptions, toward models that learn directly from experimental data while maintaining an autonomous, generative structure capable of simulating synthetic trials and perturbations.

Reservoir computing (RC) provides a promising solution to this modeling challenge [17, 18]. In RC architectures, a fixed, high-dimensional recurrent network (the reservoir) projects inputs into a rich internal state space, from which task-relevant outputs are extracted via linear readout [19, 20, 21]. This decoupling of dynamics and learning allows RC systems to efficiently capture complex nonlinear temporal dependencies without requiring gradient-based training. Recent developments have expanded RC applications across diverse domains, demonstrating its ability to model complex biological and physical systems [22, 18, 23], decode spatiotemporal neural activity and infer latent structures [24], reconstruct unobserved variables in chaotic systems [25], and even implement closed-loop feedback controllers for brain stimulation [26]. Moreover, RC models have been shown to reconstruct dynamical characteristics more precisely than direct computations based purely on training data [27]. Despite these advances, the application of reservoir computing to build biologically grounded digital twins of cortical dynamics, trained directly on multi-electrode recordings, remains underexplored. Most prior RC applications in neuroscience have focused on signal decoding or system identification rather than constructing autonomous, trial-by-trial models of cortical computation capable of virtual experimentation.

Here, we introduce a digital twin of the primate PMd based on multi-unit activity (MUA) recorded during a stop-signal reaching task, modeled through a reservoir computing framework. Our digital twin captures the low-dimensional latent dynamics underlying motor preparation, execution, and inhibition, generalizes to untrained behavioral conditions, and allows inference of latent quantities such as stop-signal reaction time (SSRT). Through systematic virtual experiments and perturbations of initial states, we reveal how internal geometry, trial-to-trial variability, and noise modulate motor behavior. This work establishes reservoir computing as a novel and powerful tool for moving from observational descriptions of cortical activity toward mechanistic, generative modeling of decision dynamics in the primate brain.

## Results

The neuronal correlates of motor planning and inhibition were investigated in two macaque monkeys (P and C) trained to perform a stop-signal reaching task [11, 6]. In this task, the subjects touched a central cue on the screen and waited for a “Go” signal, which consisted of the appearance of a peripheral target accompanied by the disappearance of the cue (Fig. 1a). In no-stop trials, after a reaction time (RT), the monkeys initiated an arm movement to reach the target (“Mov. On”). In 25% of randomly selected trials, the central cue reappeared, signaling the monkeys to withhold movement. This “Stop” signal occurred after a stop-signal delay (SSD) randomly sampled from a distribution adjusted using a staircase protocol to induce approximately 50% erroneous responses [11, 6]. In these cases, trials with overt arm movements were classified as wrong-stop trials, whereas those with successfully inhibited planned movements were classified as correct-stop trials.

**Figure 1.**
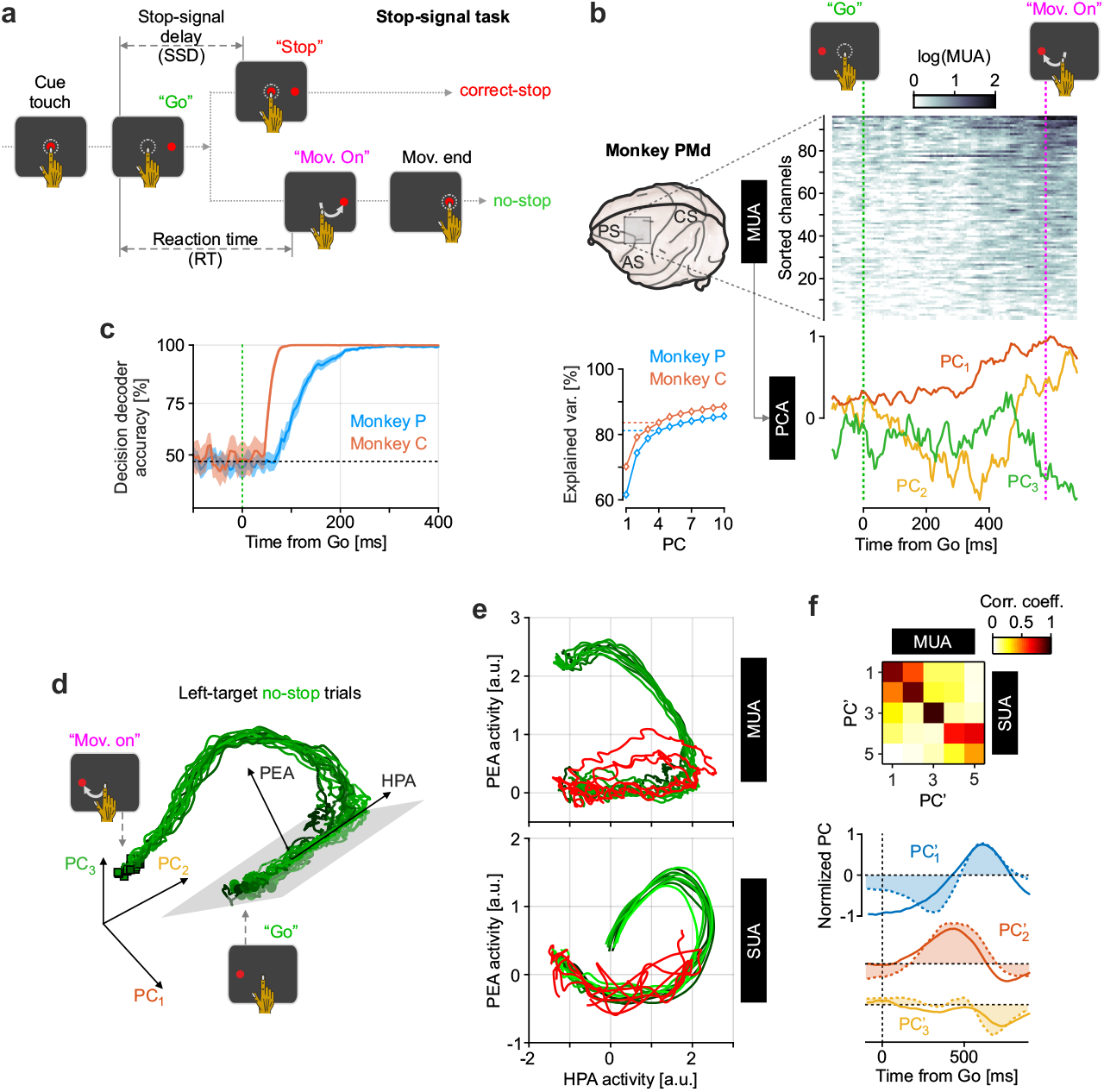
Low-dimensional MUA trajectories of motor inhibition in PMd. **a**, Stop-signal reaching task. Monkeys were instructed to reach a peripheral target (no-stop trials) after the “Go” signal (cue disappearance), while in a subset (25%) of trials a “Stop” signal (cue reappearance) presented at an unpredictable stop-signal delay (SSD) required them to withhold the arm movement. “Mov. On”, movement onset; “Mov. End”, movement end; RT, reaction time (see Methods for details). **b**, Top-left: recording sites in the left dorsal premotor cortex (PMd) recorded by intracortical multi-electrode arrays (MEA, 96 channels) in two macaque monkeys. AS, arcuate sulcus; PS, principal sulcus; CS, central sulcus. Top-right: raster plot of multi-unit activity (MUA) for an example no-stop trial during the reaction time, from the “Go” signal to “Mov. On”. MEA channels are sorted by mean MUA during the RT. Bottom-right: first three principal components (PCs) of MUAs in the same trial. Bottom-left: variance explained by the first 10 PCs derived from the PC analysis (PCA) performed around RTs and the SSDs of all no-stop and stop trials (see Methods for details). **c**, Decoding accuracy (area under the ROC curve) of motor intentions (left vs. right reach) in no-stop trials using a linear decoder during the RT for both monkeys. **d**, Neuronal (MUA) trajectories in the space of the first three PCs for no-stop trials during RTs when reaching toward the left target. Trajectories are grouped and averaged by RT. Darker (lighter) trajectories correspond to slower (faster) RTs. Holding and Planning Axis (HPA) and Planning and Execution Axis (PEA) were computed as in [6]. **e** Projection of MUA (top) and single-unit activity (SUA, bottom) trajectories for no-stop (green, from “Go” to “Mov. On”) and correct-stop trials (red, from “Go” up to 300 ms after “Stop”) onto the (HPA,PEA) plane. Bottom SUA panel adapted from [6]. **f**, Top: correlation matrix between PCs (PC’_*n*_) of average MUA and SUA profiles across channels during RTs in all reaching directions (left and right). Bottom: first three PC’_*n*_ for MUAs (solid lines) and SUAs (dashed lines with shaded areas).

After a training period, electrophysiological activity was recorded from the contralateral PMd of the employed arm using 96-channel intracortical multi-electrode arrays (MEA) chronically implanted in both monkeys (Fig. 1b, left). For each MEA channel, multi-unit activity (MUA) was extracted as the high-frequency spectral power of the raw (unfiltered) recorded signal ([28], Materials and Methods). Similar to single-unit activity (SUA), MUAs in motor cortex and PMd have been shown to encode both motor plans [29, 30, 12] (Fig. 1c) and movement inhibition [12]. However, compared to SUAs, MUAs offer the advantage of enabling decoding at both single-channel and single-trial level, thereby facilitating the design of effective brain-machine interface application [31].

### Low-dimensional MUA trajectories encode motor planning and inhibition in PMd

At the population level, single-unit activity in motor and premotor cortices is now known to be highly coordinated, generating stereotyped, action-dependent trajectories within low-dimensional neural subspaces [32, 4, 33, 8]. This theoretical framework proposes that both actions and motor plans are determined by the initial state of the cortical network, which behaves as a quasi-deterministic low-dimensional nonlinear system. Overt movements emerge only when neural trajectories exit a null-potent subspace [15, 8]. Importantly, this principle applies not only to the encoding of motor planning and execution, but also to movement inhibition and plan cancellation [6]. Does the same picture emerge when examining MUA in the PMd of monkeys engaged in a stop-signal reaching task?

Here we address this question by inspecting the spatiotemporal organization of MUAs at the single-trial level during the reaction time (Fig. 1b, top-right) and performing principal component analysis (PCA) (Fig. 1b, bottom-right; Materials and Methods). Similar to SUAs in PMd, only a few principal components (PCs) are required to consistently capture the temporal evolution of MUA across trials. In both monkeys, just the first four PCs were sufficient to explain at least 80% of the variance (Fig. 1b, bottom-left).

The MUA extracted from our PMd recordings revealed reproducible trajectories in the subspace of the top three PCs across trials. In no-stop trials, trajectories grouped and averaged by RT (Materials and Methods) transitioned smoothly from the “Go” to movement onset, resulting in the characteristic arc-shaped patterns previously observed for SUA in premotor cortex during reaching tasks [34, 8] (Fig. 1d). Remarkably, trajectories with different RTs were so stereotyped that they formed a narrow bundle, suggesting a deterministic force field that drives the neural state toward nearly identical reaching movements. This is consistent with findings from SUA recordings in monkey premotor cortex, where only initial conditions have some predictive power for the forthcoming RTs [33, 35, 36].

Following [6], we computed the “holding and planning” axis (HPA) and the “planning and execution” axis (PEA), which geometrically separate successful inhibitions from movement-generating states in SUA. The PEA defines the orthogonal (gray) plane in which neural states drift, immediately after the “Go” signal, along the HPA direction (see Materials and Methods). The initial drift in neural activity reflects a canonical ramp-to-threshold profile associated with the accumulation of evidences necessary for motor plan maturation, leading to the decodable separation of trajectories associated with left and right targets. When projected onto the (HPA, PEA) plane, both MUA and SUA followed highly similar, condition-specific trajectories (Fig. 1e). Notably, correct-stop trials (red) in both activity types remained restricted to the HPA (i.e, PEA *≃* 0), while no-stop trials (green) evolved coherently along the PEA, supporting a functional dissociation between movement execution and suppression within the PMd neural state space.

To further demonstrate the equivalence of the information encoded by MUAs and SUAs in our experimental setting, we performed a PCA on the average activity profiles for each channel/unit and condition within the interval [− 0.1, 0.9] s around the “Go” signal. We then compared the leading PCs for MUA and SUA and found a remarkably high correlation (*ρ >* 0.7) between PCs of the same rank (Fig. 1f, top; 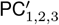). This similarity was even more apparent when matching the top three PCs, highlighting that the spatiotemporal profiles of SUA and MUA are nearly indistinguishable (Fig. 1f, bottom).

MUAs in PMd of monkey performing a stop-signal reaching task quantitatively preserve the low-dimensional geometry observed in SUAs, which underlies both motor decision and movement inhibition. Within this framework, a threshold along the PEA can be identified that must not be crossed to successfully withhold arm movement upon perceiving a “Stop” signal. According to [6] for SUAs, this threshold defines the boundary of an “output-null” region, where neural activity is confined without generating overt movements.

### RNN-based digital-twins faithfully replicate PMd MUAs

During motor planning, movement execution, and cancellation, PMd MUAs behave as a low-dimensional dynamical system endowed with a certain degree of stochasticity. The existence of an output-null region in the latent neural state space, with a threshold that must be crossed to initiate an arm movement, suggests an underlying nonlinear attractor dynamics capable of influencing behavioral output. Here, we aim to uncover these dynamical mechanisms and elucidate how they govern both the timing and the process of movement production, as well as its inhibition. To this end, we infer recurrent neural networks (RNNs) that replicate the measured MUAs and are capable to acting as a “digital twin”, that is, a generative model producing new data samples statistically indistinguishable from the experimental recordings [37, 18].

According to the reservoir computing approach [20, 19, 21], we designed a RNN composed of *N* = 2000 units that nonlinearly amplify the synaptic input, determined by a random and fixed connectivity matrix (Fig. 1a; see Materials and Methods). The RNN (i.e., the reservoir) transforms the input sequence comprised in our case by the top 4 PCs of the PMd MUAs, into a high-dimensional, nonlinear representation. Similar to an “echo-state” network [19], the same PCs are read out from the RNN’s internal state 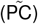 by fitting the output weights using simple linear regression on the training set. This set consists of the time series around the RTs of the first 50 no-stop trials, and from the “Go” to 300 ms after the “Stop” of the first 40 (20 for Monkey C) correct-stop trials.

Hyperparameter optimization revealed that the best-performing networks were those with random Gaussian weights having standard deviation (SD) *g* = 0.8 and *g*_in_ = 0.15 for recurrent and input connections, respectively (Suppl. Fig. 1). The heterogeneity of the relaxation time constants of the RNN units also emerged to be a crucial factor. Indeed, a uniform distribution of time constants led to substantially better performance than fixed or narrowly distributed values, enriching the RNN’s capacity to integrate information over multiple time scales. As a result, the quality of the reproduction of experimental PCs was remarkable both in the training phase (Fig. 1b) and in the test set (i.e., the trials not used during training), where the residuals 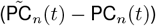 displayed a quasi-Gaussian distribution with SD much smaller than the range spanned by experimental PCs (Fig. 1c-left).

With these results, we replaced the experimental PCs used as input with the readouts 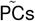, to which we added independent Gaussian white noise with the same standard deviation as the corresponding residuals (Fig. 1c-right). With this approach, we aimed to build an autonomous RNN that, at each time step, received an input statistically equivalent to the PCs of PMd MUAs recorded in each monkey. In this way, the autonomous RNN was identical to the pre-learning one but featured an updated synaptic matrix, obtained by adding the rank-1 matrices given by the outer products between the input weight vectors (fixed) and the output ones (learned). By adding noise sources, we also turned the autonomous RNN into a stochastic dynamical system, whose units were driven by correlated, memoryless Gaussian fluctuations (see Materials and Methods).

To test the capability of this RNN to operate as a digital twin, we drove it with experimental PCs in an open-loop configuration (as in Fig. 2a) until the “Go” signal, after which we let it evolve autonomously (as in Fig. 2c), generating, for each no-stop trial, new data samples, that is, the *in silico* 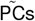. From the internal RNN state, we also read out the timing of the “digital” movement onset (i.e., the digital RTs of single trials). In the raster plots shown in Fig. 2d, we then sorted the no-stop trials according to the experimental RTs (left) and the *in silico* ones (right), obtaining 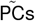 that closely mirrored those observed in experimental recordings. Both fast and slow RT trials preserved distinct activation profiles, indicating that the model captured not only the stereotyped movement dynamics but also the trial-by-trial variability inherent to cortical activity. Interestingly, also the behavioral output of the monkeys was faithfully reproduced, as evidenced by the remarkable overlap of RT distributions (Fig. 2d). Note that, although statistically equivalent, the experimental PCs in a given trial did not necessarily match quantitatively the *in silico* 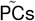 for the same trial. Indeed, the added noise played an active role in driving the exploration of internal states, effectively acting as an “energy” source (similar to [38]) that enabled the RNN to unfold a richer dynamical repertoire, without scrambling the neural trajectories, which–when grouped and average by RT–visited a manifold very close to the average experimental trajectory (Fig. 2f).

**Figure 2.**
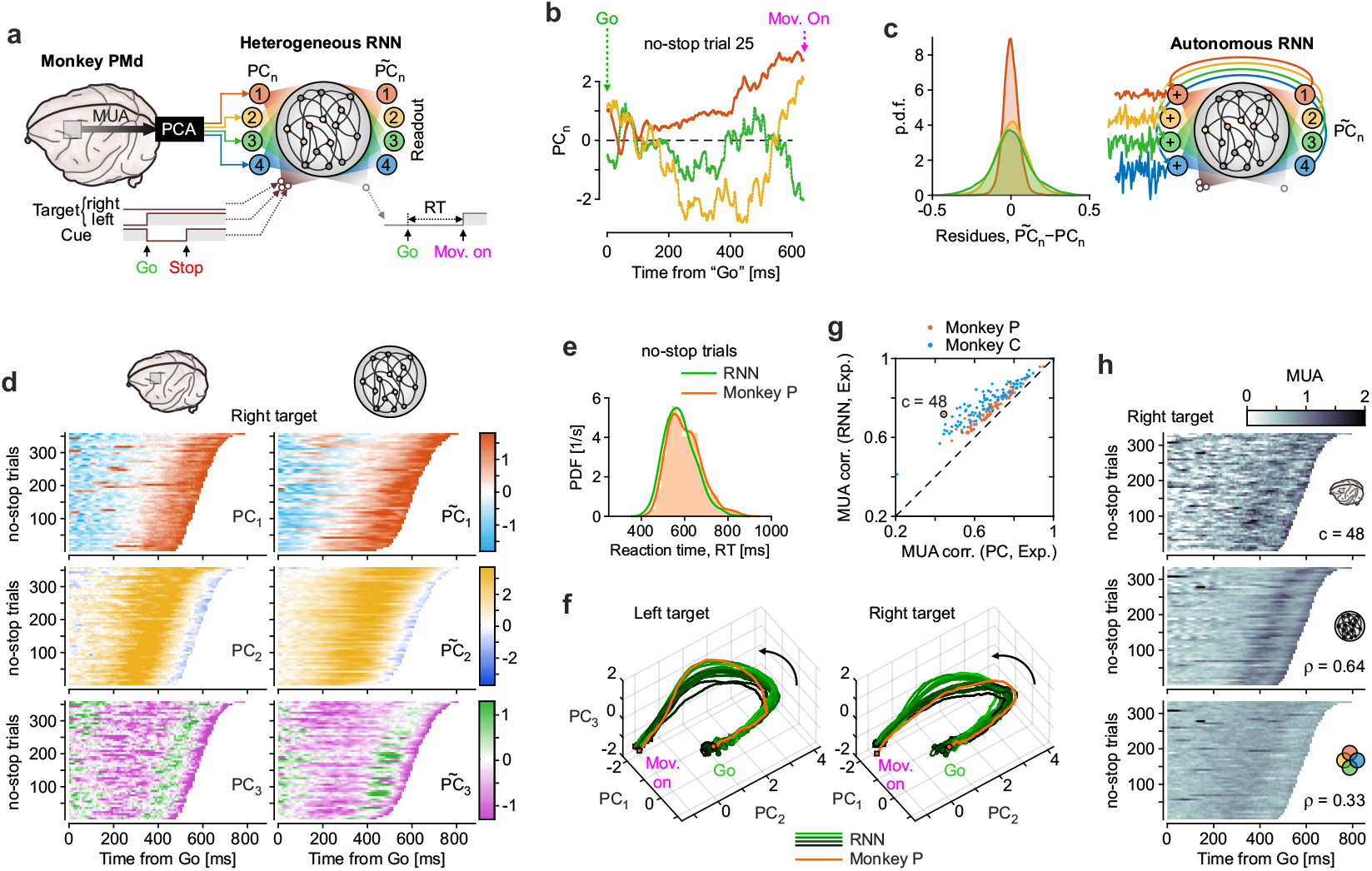
RNN replicas of PMd MUAs at single-trial level. **a**, RNN used to replicate the first four PCs obtained from the PMd MUAs, as in Fig. 1. Each network consists of *N* = 2000 units coupled through random, fixed Gaussian synaptic weights with zero mean and standard deviation (SD) *g* = 0.8. PC time series and external task-related signals (cue, target onset, and offset) are provide as input trough seven independent channels, each weighted by random, fixed Gaussian connections with zero mean and SD *g*_in_ = 0.15 (left shades). Relaxation time constants of network units are randomly sampled from a uniform distribution in the range [0.005, 0.1] s. As in echo-state networks, linear readouts (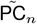, right shades) of the internal RNN state were inferred to reproduce the input PCs using ridge regression. The training set for these output weights consisted of the first 10% no-stop trials and one-third of correct-stop trials. One of the readouts was a stepwise variable representing the movement onset. Parameter values were empirically found to optimize the performance of the inferred RNNs (see Materials and Methods). **b**, First three experimental PCs (solid lines) from the same no-stop trial in Fig. 1b, with the corresponding 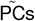 generated by the inferred RNN (dots). **c**, Left: distribution of the residues 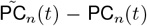 across all no-stop trials in Monkey P (*n* 3). Right: schematic of the inferred RNN in closed-loop (autonomous) configuration, where the readouts 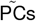 are fed back to the network, replacing the experimental inputs, with added white noise with means and SDs matching those of the measured residues. **d**, Raster plot of the first three PCs (left, Monkey P) and the corresponding 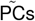 read out from the autonomous RNN (right). No-stop trials with right target sorted by experimental RTs and by the RT read out from the RNN, respectively. Autonomous RNNs are driven by experimental PCs until the “Go” and then allowed to evolve freely until the end of the “digital” RT. **e**, Left and right no-stop RT distribution in Monkey P (green) and in the autonomous RNN (orange). **f**, No-stop neural trajectories of the autonomous RNN in the top three PC subspace grouped and sorted as in Fig. 1d (green), along with the average PMD MUA trajectories (orange) for both left and right targets. **g**, Channel-wise correlation between experimental MUAs and i) MUAs reconstructed from PCs provided as input to the inferred RNN (x-axis), and ii) MUAs read out from the PC-driven inferred RNN for both monkeys. **h**, Same raster plots as in (d) for the MUA of the channel highlighted in (g) but only for test trials. Experimental data: top. Readout from inferred RNN: center. Reconstructed from the top four PCs: bottom.

Here, it is important to note that the high performance of our digital twin was achieved the information provided by PMd activity being far from complete. Only few top PCs were considered, and the chronic MEAs recorded MUAs from just a fraction of cortical and subcortical networks involved in execution of the stop-signal task. This implies that the inferred dynamical primitives effectively uncovered and incorporated such missing information. This is something expected for this class of RNNs, which possess the so-called “echo state property” [19, 39], allowing them to recover missing information through a form of dimensional embedding [40, 41, 42]. Indeed, echo-state networks can infer internal state variables of a low-dimensional and determinist dynamical system even when these variables are not directly provided as input [25]. We then tested this hypothesis in the inferred RNNs by training them to read out, from their internal states, not only the four experimental PCs but also all the MUAs recorded in the PMd of the two monkeys. The fidelity of the reconstruction was assessed by correlating the experimental MUAs with those read out from the inferred RNN in open-loop configuration (Fig. 2g). Pearson correlation coefficients were computed channel by channel across the test set. The resulting distribution of correlations was strongly skewed toward high values, with a median coefficient exceeding 0.8. Notably, the MUAs reconstructed directly from the four experimental PCs never showed a higher correlation with the actual MUAs than those with the ones read out from the RNN itself, a difference clearly illustrated by the paradigmatic example shown in Fig. 2h.

Together, these results demonstrate that the inferred digital twin not only reconstructed the observable PMd activity, but also uncovered hidden components of its latent dynamics embedded within the high-dimensional RNN state space. Finally, the introduction of an untuned, memoryless stochastic component into its units enabled us to determine of a generative model that faithfully capture the inter-trial variability observed in both neural activity and behavioral output.

### Early PMd readiness underlies reaction time variability

The ability to rapidly respond to salient environmental cues is critical for successful performance in the stop-signal task. To elucidate the neuronal basis of reaction time variability to the “Go” signal, we analyzed the inferred RNN states as a proxy for the internal PMd dynamics in each monkey.

For each no-stop trial performed by both monkeys, projecting PMd MUAs onto the HPA (i.e., the axis where neural trajectories drifted at the beginning of the reaction time) can reveal some order in the period preceding the “Go” signal. The ordering of the pre-Go HPA activity became more evident when trials were sorted by the mean digital RTs (RT_RNN_s) generated by the autonomous RNN (Fig. 3a). Specifically, for each no-stop trial, the inferred RNNs in Fig. 2 were driven by the MUA PCs until the “Go,” after which they evolved autonomously across 500 independent noise realizations. This procedure generated a distribution of RT_RNN_ for each initial condition (i.e., each experimental no-stop trial; Fig. 3b). Interestingly, some initial states yielded consistently faster or slower mean RTs, despite identical network structure and task conditions. Such variability in the mean RT_RNN_ correlated significantly with empirical RTs (Fig. 3c) for both monkeys. This finding confirmed that the inferred digital twins captured a plausible structure of behavioral variability and highlighted that the RT distribution was governed by two distinct sources of stochasticity. The former pertained to the “thermal” endogenous noise incorporated in the RNNs, likely representing fast fluctuations of neuronal activity [43, 44, 45]. The second arose from the RNN state at “Go” (i.e., the initial condition), which was determined by the pre-Go PMd activity, implying that a latent pre-Go geometry was a dominant source of behavioral heterogeneity.

**Figure 3.**
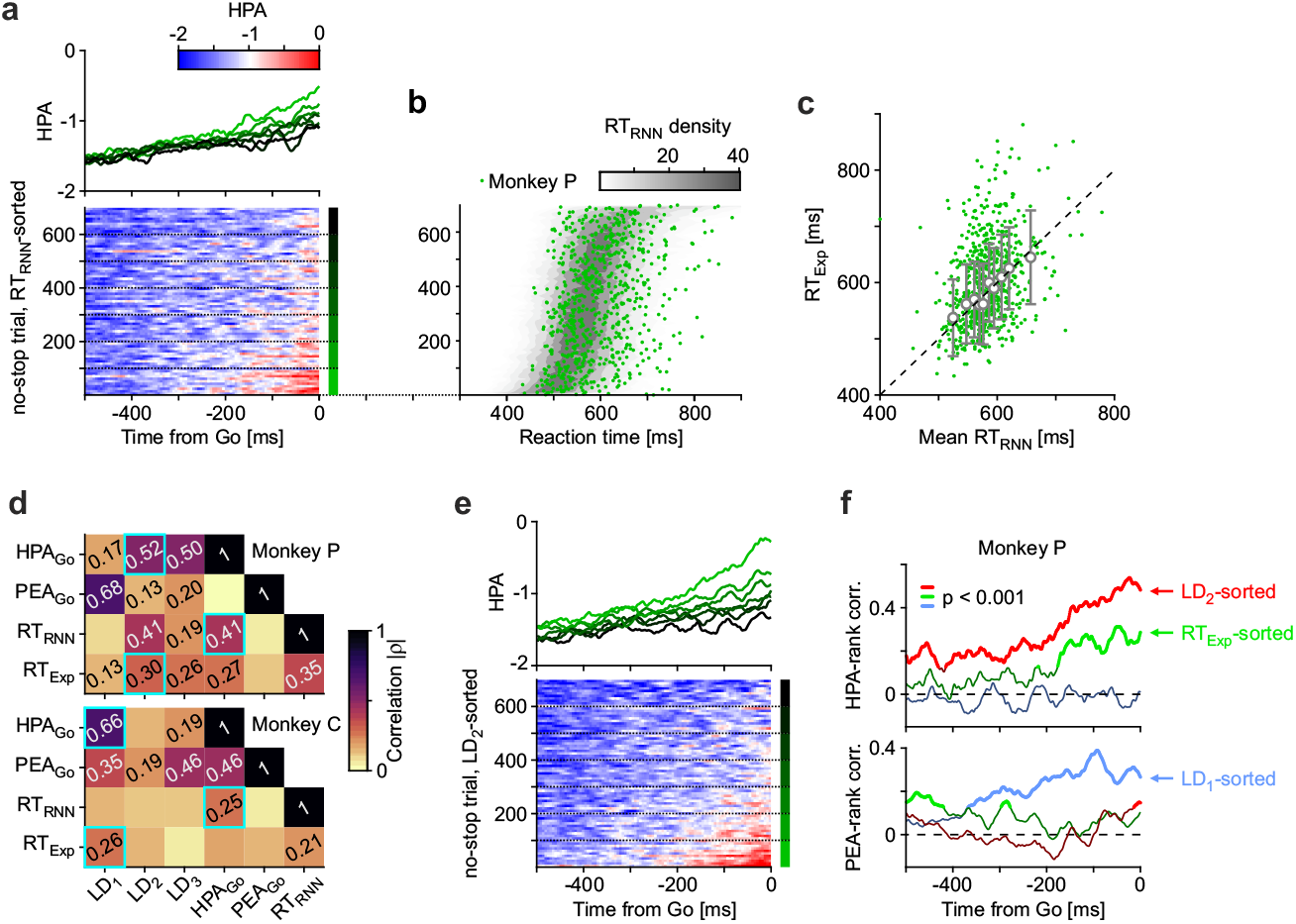
Inferred RNNs reveal how pre-Go PMd activity predicts reaction time. **a**, PMd activity in Monkey P projected onto the related HPA during the 500 ms preceding the “Go” signal across all no-stop trials (left and right targets combined). Bottom: raster plot of HPA activity sorted by the mean digital RTs (RT_RNN_s) generated by the autonomous RNN (Fig. 2c-f), shown in (b). Top: mean HPA activity for seven equally sized trial groups (indicated by the bottom green bars). **b**, Distribution of RTs (gray shading) generated by the autonomous RNN. The inferred RNN driven by MUA PCs until “Go,” evolves autonomously 500 times for each trial with different realizations of the endogenous noise. The mean RT across the resulting 500 RT_RNN_ is used to sort the experimental no-stop trials. Green dots indicate monkey RTs (RT_Exp_) for the corresponding sorted trials. **c**, Correlation between mean RT_RNN_ and monkey RTs (each green dot represents a trial). Gray circles show mean experimental RTs for each decile of mean RT_RNN_ with related SDs (error bars). **d**, Absolute Pearson correlation (| *ρ*|) across trials for both monkeys between HPA and PEA activity at “Go”, experimental and mean digital RTs, and RNN activity at “Go” projected onto the top three principal components (LD_*n*_). Cyan boxes highlight maximum *ρ* values for HPA_Go_, RT_RNN_ and RT_Exp_. **e**, Same as in (a) with trials sorted by LD_2_ of the RNN at “Go”. **f**, Correlations *ρ*(*t*) during the pre-Go 500 ms between HPA (top) and PEA (bottom) with trial rank resulting from sorting by LD_1_ (blue), LD_2_ (red) and RT_Exp_ (green). Colored dots indicate time points where |*ρ*| values are significant (*p <* 0.001).

We searched for such geometrical organization in the latent state space embedded within the inferred RNN. To this end, we performed a principal component analysis of the internal RNN at the “Go” in each no-stop trial. We found that one of the projections onto the first top PCs (LD_2_ for Monkey P and LD_1_ for Monkey C) exhibited the highest correlation with the experimental RTs (Fig. 3d). We then sorted trials by LD_1_, LD_2_ and experimental RT (RT_Exp_) (Fig. 3e,f), which revealed a more pronounced separation of HPA activity that persisted for at least 500 ms before the “Go” signal. Specifically, correlations during the pre-Go period between LD_2_ (LD_1_ in Monkey C) and HPA activity were the highest and consistently significant (*p <* 0.001). These results indicate that trial-by-trial differences in RTs were not random but structured by a low-dimensional subspace of the RNN’s internal activity, representing a latent axis of PMd readiness encoded in cortical activity well before the onset of the “Go” signal.

The digital twin thus reveals how behavioral variability in reaction time emerges from a deterministic structure within the high-dimensional cortical state space, shaped by noise, history, and intrinsic preparation. This findings further advance previews results on the neural basis of RT variability [33, 35, 36, 46], highlighting the power of digital twins to capture complex brain-behavior relationships.

### PMd readiness modulates movement inhibition

To probe the motor inhibitory capacity of the digital twin, we systematically evaluated its behavior under simulated stop-signal conditions, extending beyond training regimes. Our central aim was to determine whether the inferred RNNs, trained solely on no-stop and correct-stop trials, could autonomously reproduce inhibition-related behavior observed in primate experiments, including the emergence of realistic stop-signal reaction times (SSRTs).

One of our main findings is that this is indeed the case: the inferred RNNs autonomously generated stop trials that erroneously resulted in an “overt” movement (Fig. 4a). As expected in the stop-signal task, such errors occurred when the stop-signal delay (SSD) was too long [9]. Moreover, neural trajectories of wrong-stop trials closely overlapped with no-stop trajectories observed experimentally (compare Fig. 4b with Fig. 1e), consistent with previous analyses of the latent state space of PMd single-unit activities [6]. The same RNNs were presented with an additional set of SSDs never used in experiment. Even in this case, the resulting neural trajectories in correct-stop trials unfolded coherently with those derived the PCA of PMd MUAs (compare the red curves in Fig. 4b with those in Fig. 1e). Even more interestingly, successful stop trials were those in which PEA activity was maintained below a threshold value, as previously reported in [6]. Notably, correct-stop trajectories exhibited a clear change of direction following the “Stop” signal tending to return toward the starting point when the peripheral target appeared on the screen, signaling “Go” (Fig. 4a). The “Stop” signal then operates in PMd by rendering the initial state at “Go” attractive, thereby inducing an attractor dynamics of MUAs that eventually relax to the null-potent state space after the imperative command to cancel the movement as been successfully processed. The modulation of the force field does not affect the state space above the PEA threshold, where, even in the presence of the “Stop” signal, wrong-stop trajectories unfold unperturbed, proceeding toward the “Mov. onset” state just as in no-stop trials.

**Figure 4.**
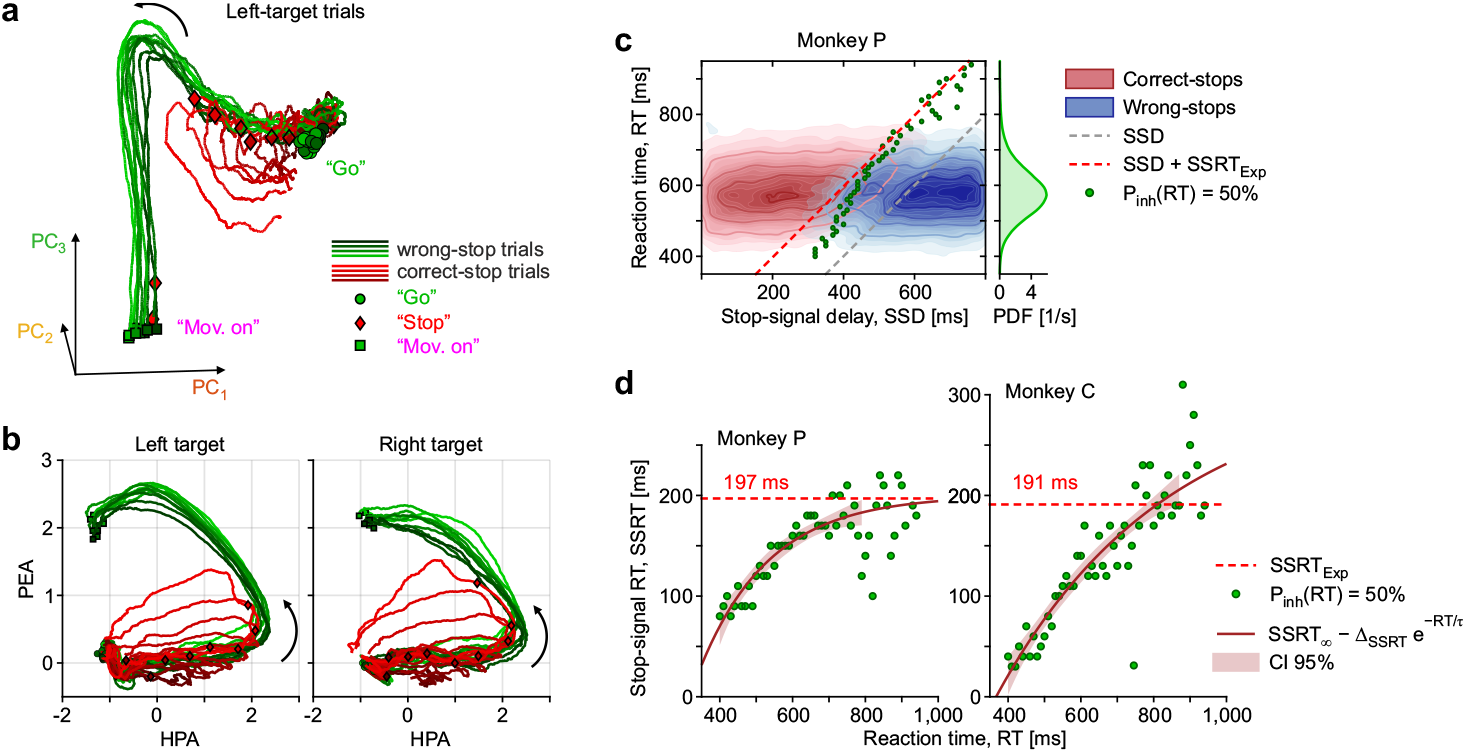
Digital-twin stop-trials uncover readiness-dependent inhibition. **a**, Neural trajectories as in Fig. 1d, generated by the autonomous RNN inferred from PMd MUAs of Monkey P. Initial conditions for target direction were selected from the the 10 no-stop trials with mean RT_RNN_ closest to the experimental mean RT. For each of these trials, 100 trajectories were generated with different and independent endogenous noise, presenting the “Stop” signal at random stop-signal delays (SSDs) between 0 and 800 ms. Green trajectories represent wrong-stop trials averaged into 10 groups sorted by RT_RNN_. Red trajectories represent correct-stop trials averaged in 10 groups sorted by SSD. Darker trajectories correspond to longer RTs and shorter SSDs, respectively. Red diamonds indicate mean SSDs in the grouped trials. **b**, Same digital trajectories projected onto the (HPA, PEA) plane for trials with left and right targets. **c**, Same simulations as in (a) and (b), with initial conditions determined by driving the inferred RNN to the “Go” for all no-stop trials. For each trials, 500 digital trajectories were generated by uniformly sampling the SSD in the [0, 800] ms-range (see Materials and Methods). Red and blue shadings represent the density of correct- and wrong-stop trajectories, respectively. For each initial condition (sorted here by mean RT_RNN_), the inhibition function *P*_inh_(*RT*) was computed; green circles correspond to *P*_inh_(*RT*) = 50%. Gray and dashed lines indicate SSD and experimental SSRT added to SSD, respectively. **d**, SSRTs estimated as the distance of the green circles from the SSD at varying RT_RNN_, for both Monkey P and C. Dashed lines: the SSRTs estimated from the experimental inhibition function. Solid lines: nonlinear fit with an exponential adaptation function with its 95% confidence interval shown as shading.

Having the ability to generate PMD-like neural trajectories under unprecedented task conditions, we were able to implement an inhibition function protocol in which the inferred RNN was stimulated from a fixed initial state across a range of SSDs. This enabled the estimation of the ‘invisible’ SSRT using standard behavioral metrics: the SSD yielding 50% stop failures was subtracted from the average RT of the corresponding no-stop simulations. However, inhibition functions often deviated from monotonicity due to undersampling at SSD extremes, and SSRT estimates were highly sensitive to rare or ambiguous trials, limiting the robustness of this approach [9, 47, 48]. For this reason, we designed an *in silico* experiment that closely mimicked the structure of experimental sessions but densely sampling also extreme SSDs. For each monkey, the inferred RNN was initialized using the pre-Go dynamics of all recorded correct- and wrong-stop trials, and then presented for 500 times “Stop” signals with SSDs uniformly drawn from the range [0, 800] ms. In this simulation framework, the inferred RNN evolved autonomously after the “Go” signal, producing neural trajectories from which decode the timing of movement onset, if any. In Fig. 4c the distribution of resulting correct- and wrong trials are shown after sorting the initial condition according to the mean RT_RNN_, as in Fig. 3b. We then estimated the SSRTs with the above integration method, grouping sorted trials and averaging them (green circles in Fig. 4c; see Materials and Methods). As a result, a clear modulation of SSRTs emerged in the digital twins of both monkeys (Fig. 4d). This modulation was well fitted by an exponential adaptation function (SSRT_∞_ −Δ_SSRT_*e*^−*RT/τ*^), approaching and crossing the mean SSRT estimated in the experiments (197 ms and 191 ms for Monkey P and C, respectively). Mean RTs then correlated with SSRTs indicating that the readiness of the PMd similarly influence the response to both the “Go” and “Stop” signals, in some agreement with the hypothesis that the go and stop processes in the race model [9] are not independent [49].

The developed digital-twin approach thus proved to be effective in providing a mechanistic bridge between latent cortical dynamics and observable behavioral control, enabling the inference of covert quantities such as trial-specific SSRTs and the contextual flexibility of motor inhibition.

## Discussion

This study introduces a biologically grounded digital twin of the PMd, based on a reservoir computing architecture trained directly on MUA recorded during a stop-signal task. By embedding MUA-derived latent trajectories into a recurrent dynamical system, the model not only replicates the empirical structure of PMd activity but autonomously generates behaviorally interpretable outputs, enabling direct causal interrogation of cortical computation. Our results demonstrate that MUA, despite lacking the single-neuron resolution of SUA, contains sufficient information to reconstruct well-defined motor trajectories in low-dimensional state space. The trajectories extracted from MUA data accurately recapitulated known PMd dynamics, including preparatory divergence along the HPA and movement-aligned evolution along the PEA, in line with prior findings based on SUA and latent dynamics analysis [6, 15, 36, 8]. Importantly, the digital twin learned to generalize these latent structures beyond the training trials, exhibiting high-fidelity MUA reconstructions and improved performance compared to direct PCA-based decoders. The superiority of the inferred RNN depended critically on its internal architecture. Networks with uniform distributions of neuronal time constants *τ*, optimal spectral radius of the recurrent synaptic matrix, and moderately injected internal noise exhibited rich yet stable dynamics capable of capturing the full temporal structure of PMd trajectories. This confirms previous theoretical work demonstrating that heterogeneous timescales and controlled instability promote flexible memory and trajectory embedding in recurrent systems [50, 27, 51].

The RNNs we inferred from the PMd MUAs, implemented an echo state network as only the readout weights need to be learned, which reduces the number of parameters to fit. This is an important advantage when the dataset is limited in size, exploiting the theoretical framework of reservoir computing, where RNNs serve as universal approximators of dynamical systems [52, 53, 54], effectively capturing low-dimensional and stochastic dynamics. We refined the initially random connectivity matrix by adding a low-rank update obtained through linear regression to read out the first four PCs of PMd MUAs. This low-complexity feedback mechanism further facilitated systematic dissection of the analog computation implemented by the RNN dynamics [55, 56]. This kind of RNN-based digital twins enabled detailed inspection of the underlying dynamical system, allowing the application of “opening the black box” methods to interpret RNN computations [57].

Previous work has rarely demonstrated RNNs replicating MUA time series directly; instead, more sophisticated hybrid models combining RNNs with variational auto-encoders have been proposed [58]. While RNNs have been used to replicate trial-averaged neural activity profiles by updating all recurrent synaptic weights [59], such approaches are not generative in the sense of sampling novel trajectories. More specifically, within the motor decision-making framework, RNNs have largely been employed to replicate muscle activity [60, 14] rather than cortical population dynamics. To our knowledge, our results are among the first to prove the effectiveness of ESN to build a digital twin that generates and reproduces the spatiotemporal structure of cortical MUA, enabling a generative, data-driven model of cortical population activity.

In the study of movement inhibition, behavioral performance in the stop-signal task is classically described by the race model [9]. This model conceptualizes response inhibition as a race between two independent processes: a go process that initiates a response and a stop process that inhibits it. The outcome depends on which process finishes first, with the latency of the stop process (i.e., the SSRT), estimated indirectly. More recently, the interactive race model [49] has extended this framework by maintaining independence between go and stop processes for most of their duration but introducing a late-stage interaction that determines whether a response is ultimately executed or canceled. Using virtual experiments with the inferred RNNs on an unseen set of stop-signal delays (SSDs), we demonstrated that premotor dorsal cortex (PMd) readiness predicts both reaction time to initiate reaching movements and the velocity of canceling planned actions. This highlights a temporally distributed interaction between go and stop processes as conjectured in [6], providing novel insights into movement inhibition beyond classical models.

A central finding of our work is that trial-by-trial variation in RT arises not only from random fluctuations, but also from structured differences in the reservoir internal state at movement onset. Specifically, the projection of the Goaligned state onto a suited one-dimensional manifold robustly predicted the mean RT across hundreds of random realization of the endogenous noise. This result aligns with experimental evidence that preparatory activity in motor cortex modulates the timing of action execution [61, 33, 35, 36, 62]. In this work, the RNN-based digital twin we inferred replicated behavior consistent with motor inhibition. Using a virtual experimental design that reproduced the temporal structure of stop-signal task performed by the monkeys, we showed that the model produced realistic SSRTs and inhibition functions. These dynamics were not programmed explicitly but emerged from the reservoir internal geometry after training on no-stop and correct-stop trials.

Together, these findings support a view of PMd as a dynamical substrate where movement planning and inhibition emerge from structured interactions among internal states. Reservoir computing provides a powerful computational framework to model such dynamics from experimental data, enabling not only decoding but generation of trial-resolved neural trajectories and behavioral variability. The digital twin thus constitutes a novel methodological advance for computational neuroscience, bridging latent cortical geometry with observable behavior, and opening new opportunities for *in silico* experimentation under biologically realistic conditions.

## Methods

### Animals, behavioral task and data acquisition

Two adult male rhesus monkeys (*Macaca mulatta*; subjects P and C, 7–9.5 kg) were trained to perform a countermanding reaching task. The task structure followed the classical stop-signal paradigm, designed to probe motor preparation, movement initiation, and the ability to cancel an ongoing action. Each trial began with the monkey touching and holding a central cue displayed on a screen. During this holding period—randomly selected between 400 and 900 ms—the animal had to maintain its hand on the starting location while keeping visual fixation. This temporal variability prevents anticipatory timing strategies. At the end of the holding period, the “Go” signal occurred, consisting of the disappearance of the central cue and the simultaneous appearance of a peripheral visual target on either the left or the right side. This instructed the monkey to initiate an arm movement toward the target. The interval between the “Go” signal and the time when the monkey lifted its hand to start the movement defines the reaction time RT. To receive a reward in these no-stop trials, the movement had to begin within 1,000 ms of the “Go” signal and proceed correctly to the target.

In approximately 34% of trials, the Go instruction was unexpectedly followed by a “Stop” signal. The Stop signal consisted of the reappearance of the central cue, indicating that the monkey should cancel the movement it was preparing to make. The “Stop” signal occurred after a variable stop-signal delay (SSD), chosen on each trial from the set {400, 500, 600, 700, 800 ms}. The SSD is the critical parameter that determines the difficulty of stopping: short SSDs correspond to early “Stop” signals and therefore easier cancellations, whereas long SSDs correspond to late “Stop” signals and therefore harder cancellations. A stop trial was classified as a correct-stop if the monkey successfully withheld movement and kept the hand on the central position for an additional holding period lasting between 400 and 1,000 ms. Conversely, if the hand left the starting position after the Stop signal, the trial was classified as a wrong-stop. Only correct-stop trials were rewarded.

To maintain behavioral engagement and obtain reliable stopping statistics, the session included a classical staircase tracking procedure. Its purpose was to dynamically adjust the SSD so that the monkey’s probability of successfully withholding the movement stayed close to 50%, a performance level essential for estimating the stop-signal reaction time (SSRT), that is, the latent processing time required to inhibit an action. The staircase operated as follows: after a correct-stop trial, the SSD was increased by 100 ms, making the next stop trial harder; after a wrong-stop trial, the SSD was decreased by 100 ms, making the next stop trial easier. Iterating this rule causes the SSD to converge toward a value at which successful inhibition occurs roughly half of the time, ensuring balanced sampling of correct-stop and wrong-stop trials and enabling stable estimation of stopping performance across the session. See [6] for further details. All behavioral procedures, animal care, housing, and surgical protocols adhered to European (Directive 86/609/ECC and 2010/63/EU) and Italian regulations (D.L. 116/92 and D.L. 26/2014) on the use of non-human primates in scientific research and were approved by the Italian Ministry of Health.

### Signal extraction and recording acquisition

Neural signals were acquired from the dorsal premotor cortex (PMd) using high-density intracortical recordings, providing access to both single-neuron activity and broader multi-unit activities. In both monkeys, 96-channel Utah arrays (BlackRock Microsystems) were implanted on the cortical surface after dura opening, using the arcuate sulcus and precentral dimple as anatomical landmarks to target PMd contralateral to the reaching arm [6]. Extracellular signals from each electrode were amplified and digitized using a Tucker-Davis Technologies RZ2 system equipped with a PZ2 preamplifier, producing continuous broadband raw recordings at a sampling rate of 24.4 kHz suitable for extracting both spike-sorted single-unit activity (SUA) and multi-unit activity (MUA; see [28, 12, 6] for further details).

### Comparison across signal types

To assess whether SUAs and MUAs convey comparable information about population dynamics in PMd, we performed a direct quantitative comparison of their dimensionality and representational structure. This analysis was motivated by the need to verify that MUA—which aggregates the high-frequency activity of small neuronal populations—can reliably capture the same low-dimensional neural geometry typically extracted from spike-sorted SUA, thereby justifying its use in the main analysis.

For the MUA dataset, we considered activity from all 96 channels of the Utah array, covering a time window from 1 s before the Go signal to 900 ms after it, with a temporal resolution of 5 ms. For the SUA dataset, we included 105 well-isolated neurons recorded from the same grid and in the same sessions. SUA spike trains were smoothed with a 25 ms uniform moving-average window to obtain continuous firing-rate estimates. To enable a fair comparison, SUA data were downsampled to 5 ms resolution, and both signal types were restricted to a shared temporal window of [− 1000, 900] ms around the “Go” signal.

We applied principal component analysis (PCA) separately to the SUA and MUA matrices. For each dataset, the input to PCA was a matrix where each row represented the trial-averaged activity of a channel or neuron concatenated across time. The first principal components (PCs) were normalized independently to the interval [−1, 1] for visualization purposes. As a quantitative measure of correspondence between the representations extracted from the two signal types, we computed Pearson correlation coefficients between the first five principal components obtained from SUA and MUA. The resulting correlation matrix, shown in Fig. 1f, revealed a strong match between corresponding components, indicating that both signals capture highly similar low-dimensional temporal motifs in PMd activity.

These results demonstrate that MUA preserves the essential structure of the population dynamics revealed by SUA, validating the use of MUA for dimensionality reduction and subsequent analyses of preparatory and movement-related activity.

### Low-dimensional latent state space analysis

To examine how population activity in PMd evolved during the stop-signal task, we projected the MUA onto a latent low-dimensional space using the top PCS. The rationale for this dimensionality reduction is that cortical population responses exhibit strong temporal covariation across channels, such that a small number of components can capture most of the task-related variance while filtering out channel-specific noise. This allows us to visualize neural trajectories in a compact space and to compare population dynamics across trial types under a unified representation Fig. 1b.

For each recording session, MUA was extracted from all responsive channels (0 discarded in Monkey P, 1 discarded in Monkey C) and segmented around task-relevant events. No-stop and wrong-stop trials were aligned to the “Go” signal and included from 50 ms before Go to each trial’s movement onset. Correct-stop trials were aligned to the “Stop” signal and included from 50 ms before Go to 300 ms after the “Stop” cue. These windows encompassed the time intervals in which preparatory, movement-related, and inhibitory-related modulations were reliably expressed (see Fig. 1d for representative averaged MUA profiles across no-stop trials).

To reduce trial-by-trial variability and enhance consistent structure, trials within each condition were sorted by behaviorally relevant metrics—RT for no- and wrong-stop trials, and SSD for correct-stop trials. Sorted trials were then grouped (no-stop trials in deciles; correct- and wrong-stop trials in quartiles) and averaged within each group. This procedure produced one condition-specific matrix per group, each representing the average temporal evolution of MUA across channels.

The matrices for all trial types were then concatenated along the temporal dimension to form a single channel-by-time data matrix for PCA. This ensured that the extracted PCs reflected population-level patterns shared across conditions, rather than being driven by a specific subset of trials. PCA was applied to this aggregated matrix, yielding orthogonal components ordered by explained variance. The first few components provided a compact representation of the dominant temporal motifs in PMd activity, capturing the coordinated fluctuations across channels that underlie motor preparation, movement execution, and response inhibition. The resulting low-dimensional projections of the population activity are shown in Fig. 1d and this latent representation forms the basis for all subsequent analyses and model comparisons.

### Projection into behavioral state subspaces, HPA and PEA identification

After obtaining the low-dimensional PCA representation of MUA activity, we identified two task-relevant directions within the latent space to summarize how PMd population activity evolves during preparation and movement execution. Following the approach introduced in [6], we defined the Holding and Planning Axis (HPA) and the Planning and Execution Axis (PEA). These axes provide a compact behavioral subspace within which neural trajectories from all trial types can be compared.

To compute PEA, we first averaged PCA trajectories across groups of No-stop trials and concatenated the initial portions of these trajectories (first 300 ms after Go) in the space of the first three principal components. We then fitted a plane to these pooled points using a least-squares procedure and defined PEA as the unit normal vector to this best-fit plane, with its sign chosen such that projections increase toward movement execution.

HPA was then defined as the main ramping direction within the same plane. Using the two in-plane orthonormal basis vectors returned by the plane fit, we projected the early no-stop trajectories onto the two-dimensional basis and collected the resulting coordinates over the same time window (0–300 ms). We then fitted a line to these points by linear regression and used the resulting slope to construct a unit vector within the plane, which was taken as HPA.

The pair of unit vectors (HPA, PEA) thus obtained defines a two-dimensional behavioral subspace. All trajectories from nostop, correct-stop, and wrong-stop trials were subsequently projected onto this subspace to compare population dynamics across behavioral conditions (Fig. 1e). This projection framework provides a unified latent-space representation of preparatory, execution-related, and inhibitory population dynamics in PMd.

### Time-resolved motor decoder analysis

To assess how directional information emerges over time in PMd population activity, we implemented a time-resolved linear decoder applied directly to the instantaneous MUA. The goal of this analysis was to quantify, with millisecond resolution, how well neural population activity distinguishes left-from right-directed movements around the moment of commitment.

For each session, we defined a temporal window spanning from 50 ms before to 200 ms after the “Go” signal. At each discrete time point within this interval, we extracted the instantaneous MUA value from every electrode across all trials, forming a trial-by-channel activity matrix. Trials were then randomly partitioned into training (80%) and test (20%) sets, and this procedure was repeated 50 times to obtain robust performance estimates.

For the training set, instantaneous activity vectors were assembled into a matrix *X*_train_ ∈ ℝ^*d×n*^, where *d* is the number of channels and *n* the number of training trials. A bias term was included by appending a column of ones. Movement direction (left vs. right) was encoded as a binary label vector *y*_train_ ∈ {0, 1} ^*n*^. A linear decoder was learned by solving the least-squares problem using the Moore-Penrose pseudoinverse:

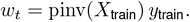

The resulting weight vector was applied to the held-out test trials to generate scalar predictions, and decoder accuracy at that time point was quantified using the area under the ROC curve (AUC). Averaging across 50 random train/test splits yielded a smooth estimate of decoding performance as a function of time.

The resulting time-resolved AUC curve (Fig. 1c) revealed a monotonic rise in directional decoding accuracy shortly after the Go cue, peaking around movement initiation. This temporal profile was consistent across sessions and monkeys, indicating that PMd population activity carries increasingly reliable movement-direction information as the neural state progresses toward execution.

### Network structure and dynamics

Reservoir computing provides a powerful framework for modeling nonlinear dynamical systems from data while keeping learning efficient and well-posed. Only the linear readout is trained, whereas the recurrent internal dynamics remain fixed, allowing the reservoir to approximate complex temporal transformations without requiring full recurrent learning, which is typically unstable or ill-posed. Echo-state networks (ESNs), the most widely used reservoir architecture [63, 19], have been successfully applied to neural population modeling, latent-state reconstruction, and forecasting [21, 18]. These properties make ESNs a suitable substrate for constructing a data-driven digital twin of PMd activity.

The reservoir consisted of *N* = 2000 units with continuous-time dynamics. Each unit *x*_*j*_(*t*) evolved according to

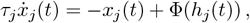

where Φ(*z*) = tanh(*z*) and intrinsic time constants *τ*_*j*_ were independently drawn from a distribution *f* (*τ*) specified in the hyperparameter search (see Section). The recurrent input was

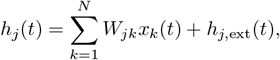

with *W* ∈ ℝ^*N×N*^ a sparse random matrix with connection probability *p* = 0.16. Non-zero synaptic weights were sampled from a Gaussian distribution with variance *g*^2^*/*(*pN*), where *g* (spectral radius) was selected through hyperparameter search. External inputs consisted of the four experimental PCs and task-event signals:

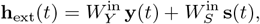

where **y**(*t*) ∈ ℝ^4^ contains the first four PCs of PMd MUA and **s**(*t*) ∈ ℝ^*S*^ encodes task events (cue, Go, target, Stop). Entries of 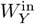 and 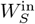 were drawn from zero-mean Gaussian distributions with standard deviations *g*_in_ and *g*_*s*_ = 0.02. Reservoir dynamics were integrated using Euler’s method with time step Δ*t* = 10^−3^. A schematic of the model appears in Fig. 2a.

During training, the reservoir was driven by the empirical PCs **y**(*t*) (teacher forcing). In this open-loop regime, internal states **x**(*t*) are uniquely determined by the input history due to the echo state property [19, 39]. Only the linear readout *W* ^out^ ∈ ℝ^4*×N*^ was optimized to reconstruct the target PCs:

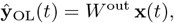

where **ŷ**_OL_(*t*) denotes the readout prediction during open-loop training, obtained while the true PCs **y**(*t*) are injected as input. The optimal readout was computed by ridge regression:

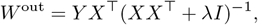

where *X* is the reservoir state matrix, *Y* contains the target PCs, and *λ* was selected by cross-validation (Fig. 2b).

After training, the model was switched to closed-loop mode to generate autonomous trajectories. In this regime, the PCs were replaced by the model’s own predictions,

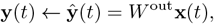

yielding the closed-loop input

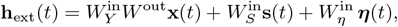

as shown in Fig. 2c. To ensure stable autonomous evolution and to reproduce the variability of PMd activity, the injected noise ***η***(*t*) was derived from the residuals of the open-loop fit. After training, we computed

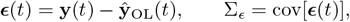

and generated noise as

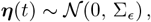

injecting it through 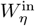, whose entries matched the distribution of other input matrices. Because the noise is scaled to empirical residual statistics, closed-loop trajectories explore realistic dynamical neighborhoods around the learned attractor and avoid convergence to trivial fixed points.

To determine whether the reservoir recovered latent structure beyond the four PCs used as input, we compared two reconstruction strategies. The first approach used the PC scores to map the four PCs directly to the MUA channels, providing the maximum variance explainable from the PCs alone. The second approach trained an identical decoder using the reservoir states as predictors. Reservoirs were trained to reproduce MUA activity over the full duration of 50 no-stop and 40 correct-stop trials in monkey P (50 no-stop and 20 correct-stop in monkey C), rather than restricting training to post-Go windows. For each channel, we computed the Pearson correlation between the measured MUA and the reconstructed signal. A Wilcoxon signed-rank test revealed significantly higher correlations for the reservoir-based decoder in both monkeys (*p <* 0.05). Moreover, the distribution of correlations obtained using the reservoir was strongly skewed toward high values (median *>* 0.8), whereas PC-based reconstructions never reached such accuracy. These results indicate that the ESN recovered latent components of PMd dynamics that were not explicitly present in the four PCA inputs, consistent with the theoretical framework of generalized synchronization in driven recurrent networks.

### Training procedure and virtual experiments

For each session and monkey, the reservoir was trained using the first 50 no-stop and 30 correct-stop trials. MUA from each trial was projected onto the top four PCs and resampled at 1 ms resolution using shape-preserving cubic interpolation. These normalized PC trajectories defined the neural input vector **y**(*t*). Three additional stepwise signals **s**(*t*)—Cue, Left-target, and Right-target—were provided to indicate the presence of the corresponding on-screen events. The full input **h**_ext_(*t*) = [**y**(*t*), **s**(*t*)] ∈ ℝ^7^ was injected into the reservoir through an input matrix *W* ^in^ whose entries were drawn from zero-mean Gaussian distributions with standard deviations *g*_in_ (neural inputs) and *g*_*s*_ = 0.02 (task-event inputs). Although the reservoir was stimulated during the entire [Start, End] period of each learning trial, only the activity within [Go, Detach] for no-stop trials and within [Go, Stop + 300 ms] for correct-stop trials was used to compute the readout weights.

The composition of the learning set followed an empirical criterion. Rather than training on a large fraction of trials, we adopted a compact 10/90 learning/testing ratio. This ensured that the training trajectories sampled the key dynamical regimes of PMd activity while limiting the risk of overfitting to specific behavioral episodes. A restricted learning set is sufficient for this system because the reservoir reproduces the neural trajectories by visiting a low-dimensional attractor manifold, and accurate mapping of this manifold does not require exhaustive sampling of all trials. During evaluation, all trials—including those used for training—were presented again in open- and closed-loop mode; because the reservoir dynamics are stochastic and each realization diverges due to noise injection, reusing training trials does not bias performance estimates.

The RNN was trained to reproduce not only the four PCs but also a binary movement variable, set to 1 at movement onset and 0 otherwise. This additional target allowed us to derive an endogenous RT from the closed-loop dynamics. After training the readout matrix *W* ^out^ ∈ ℝ^5*×N*^, the RNN was initialized by presenting the empirical trajectory over the [Start, Go] window to guarantee generalized synchronization and remove dependencies on initial conditions. After “Go”, the RNN was switched to closed-loop mode and autonomously generated both neural trajectories and the movement-output trace, from which RT was extracted by thresholding the movement output at 0.4.

This framework enabled a wide set of virtual experiments. First, we generated synthetic behavioral sessions by sampling SSDs using the same staircase rule as in the animal task, allowing us to compute trial-by-trial inhibition functions and SSRTs from hundreds of closed-loop realizations with independent noise. Second, we probed the causal role of the internal state at “Go” by testing fixed SSD values across different initial conditions, quantifying how proximity to the attractor boundary influenced the probability of successful stopping. Finally, by systematically scaling the amplitude of the stop-signal input, we evaluated how changes in the effective stopping drive altered the divergence of trajectories within the latent (HPA, PEA) plane (Fig. 4), demonstrating that inhibitory performance emerged from the RNN dynamics rather than from an explicitly coded stopping rule.

### Hyperparameters choice

The reservoir hyperparameters *g, g*_in_, *f* (*τ*), and *f*_noise_ were optimized through an extensive grid-search aimed at identifying the configuration that best reproduced the statistical structure of intracortical activity. To explore a sufficiently broad ensemble of models, for each session we generated 10 RNNs for every tested hyperparameter combination. The explored ranges were: spectral radius *g* ∈ {0.3, 0.4, 0.5, 0.6, 0.7, 0.8, 0.9} ; input scaling *g*_in_ ∈ {0.05, 0.15, 0.45, 0.75}; residual-noise factor *f*_noise_ ∈ {0, 0.25, 0.5, 1, 2, 3, 4, 5} ; and neuronal time-constant distribution *f* (*τ*), implemented either as a uniform distribution in [0.005, 0.1] s or as fixed time constants *τ* = {0.005, 0.05, 0.1} s. Each RNN was trained on the first 50 no-stop and 30 correct-stop trials and tested on all remaining no-stop trials of the same session.

For evaluation, we considered only trials in which both the experimental data and the reservoir produced a detectable RT, ensuring a consistent basis for performance comparison. Inferred RNN performance was quantified using two complementary metrics. The first metric, *z*_1_, measured the similarity between experimental and simulated trajectories for the first three PCs. For each PC, RNN trajectories were aligned by RNN-generated RTs, grouped into non-overlapping chunks (20 for Monkey C and 50 for Monkey P, reflecting dataset size), averaged within each chunk, and compared to their experimental counterparts via Pearson correlation. The final score was the average correlation across PCs and chunks,

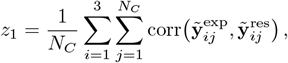

where each trajectory was truncated to its minimal common duration *L*.

The second metric, *z*_2_, quantified the discrepancy between experimental and simulated RT distributions using the log-distance between their first three central moments (mean, standard deviation, skewness), supplemented by a penalty accounting for differences in the number of detected RTs:

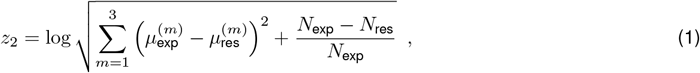

In accurately performing RNNs, the relative RT-count mismatch was typically below 1%, whereas poorly performing inferred RNNs often failed to produce any RTs.

Importantly, the joint use of *z*_1_ and *z*_2_ ensured that hyperparameter selection optimized both the fidelity of the neural dynamics and the ability of the model to reproduce behaviorally relevant outcomes. Matching only the neural trajectories or only the reaction-time distribution would have been insufficient: the first would yield models that look neurally plausible but fail to capture decision-related computations, while the second would allow correct behavioral statistics at the cost of unrealistic neural dynamics. Optimizing both metrics simultaneously was therefore essential to guarantee that the resulting reservoir behaved as a coherent dynamical system capable of reproducing, with the same parameter set, the structure of PMd population activity and the behavioral signatures emerging from it.

Grid search revealed that residual-noise amplitudes in the range *f*_noise_ ∈ [0.5, 1] consistently produced higher correlations. To select the optimal time-constant distribution, we defined an “optimal region” in the (*z*_1_, *z*_2_) plane (*z*_1_ *>* 2.5, *z*_2_ *<* − 1) and counted, for each *f* (*τ*), the number of (*g, g*_in_) pairs falling in this region. The uniform distribution of *τ* clearly outperformed all fixed values. We then restricted the search to this time-constant distribution and noise values {0.5, −1} and recomputed performance across all (*g, g*_in_) combinations. For each pair, we measured the Euclidean distance to the optimal region in the (*z*_1_, *z*_2_) plane and selected the combination with minimal distance.

The resulting optimal hyperparameters were used for all subsequent analyses and virtual experiments. They are summarized in Table 1.

**Table 1.**
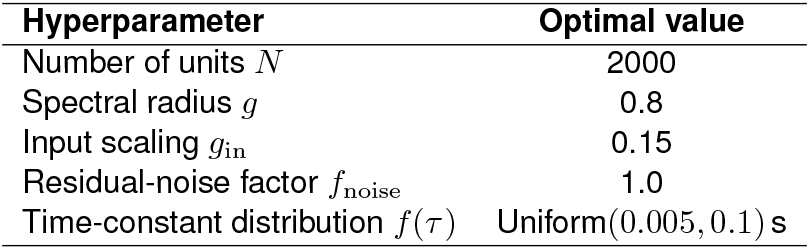
Optimal hyperparameters used in all simulations.

| Hyperparameter | Optimal value |
| --- | --- |
| Number of units $N$ | 2000 |
| Spectral radius $g$ | 0.8 |
| Input scaling $g_{\text{in}}$ | 0.15 |
| Residual-noise factor $f_{\text{noise}}$ | 1.0 |
| Time-constant distribution $f(\tau)$ | Uniform(0.005, 0.1) s |

### Reaction times dependence on initial conditions

To quantify how the internal state of the inferred RNNs at the time of the “Go” signal shapes the reaction time, we analyzed how variability in the inferred RNN’s initial conditions propagates into behavioral variability during closed-loop simulations. For each empirical no-stop trial, we computed the initial condition (IC) as the RNN state vector **x**(*t*_Go_) obtained during open-loop driving. In this phase, the RNN receives the four top PCs of MUAs as input, ensuring that **x**(*t*_Go_) is uniquely determined by the experimental history of that trial. Collecting these vectors across trials yielded a trial-by-trial set of latent initial conditions in the high-dimensional reservoir state space (Fig. 3a). These ICs provide a compact model-based estimate of the latent preparatory state from which movement-related dynamics unfold.

Because RNN states lie on a low-dimensional manifold, we reduced these vectors using PCA, retaining only the components associated with non-zero eigenvalues (Fig. 3b–c). The resulting PCs was named as latent dimensions (LD_*n*_). This ensured that the reduced representation captured meaningful dynamical structure while filtering out numerical degeneracies. The resulting embedding defined the low-dimensional space in which we compared model-generated ICs to their experimental counterparts.

To probe sensitivity to initial conditions, each empirical IC was used as a seed for closed-loop simulations. For every such seed, we ran 500 independent realizations, each with a different sample of structured residual-based noise. In all simulations, the SSD was fixed at 0 ms, allowing us to isolate the contribution of initial-state geometry and intrinsic noise on movement initiation without interference from stopping dynamics. RTs were extracted using the same movement-output threshold applied in the experimental analysis (Fig. 3c).

This procedure produced, for each empirical IC, a distribution of model-generated RTs representing the behavioral variability expected from fluctuations around that particular latent state. To compare model and experimental geometry, we repeated the same PCA-based reduction on ICs extracted directly from the experimental dataset. Model-derived ICs exhibited the same dimensionality as the empirical ones (two dimensions for both monkeys), and their spatial structure matched closely across datasets (Fig. 3d), indicating that the inferred RNN faithfully reproduces the geometry of trial-to-trial variability present at the “Go” signal.

### Evaluation of inhibitory dynamics

To characterize how the digital twin implements motor inhibition, we evaluated the inferred RNN’s response to “Stop” signals using progressively more robust analytical frameworks. Our goal was to determine whether the inferred dynamics, trained only on no-stop and correct-stop trials, could autonomously reproduce key behavioral markers of inhibitory control such as the SSRT and the inhibition function.

We first assessed the ability of the inferred RNN to discriminate between correct- and wrong-stop trials using a ROC analysis. For each experimental initial condition (IC), we generated pairs of simulated trials using identical noise realizations, one with and one without a stop signal at a given SSD. Kernel density estimates of the inferred RNN’s movement-related readout were used to quantify the separability between correct- and wrong-stop trajectories. Although this approach demonstrated partial discriminability, the inferred SSRT values were highly sensitive to both the IC and the stochastic noise realization, and the temporal alignment of ROC curves lacked a consistent behavioral grounding.

We next implemented a classical SSRT estimation based on inhibition functions. For each IC, the inferred RNN was tested across a dense grid of SSDs while keeping noise fixed. The probability of responding at each SSD was used to identify the SSD at which 50% of simulated stops failed, from which SSRT was estimated by subtracting the mean no-stop RT. While more interpretable, this procedure required a large number of simulated trials and was sensitive to irregularities and non-monotonicity in the inhibition curve, especially at SSD extremes. Moreover, variability across network instantiations revealed a strong dependence on initial-state geometry.

To overcome these limitations, we designed a virtual experimental protocol that reproduced the temporal structure of behavioral sessions. For each IC and noise realization, we generated sequences of simulated trials that interleaved no-stop, correct-stop, and wrong-stop conditions with SSDs drawn from the same staircase procedure used in animal experiments. This allowed the inferred RNN to be tested in a contextually coherent regime and enabled SSRT estimation using the standard integration method. This framework also allowed us to examine whether the inferred RNN’s internal state at “Go” predicted stopping success and to assess how inhibitory performance depended on training history.

## Acknowledgments

Work partially funded by the Italian National Recovery and Resilience Plan (PNRR), M4C2, funded by the European Union - NextGenerationEU (Project IR0000011, CUP B51E22000150006, ‘EBRAINS-Italy’) to SF and MM.

## Supplementary Material

**Supplementary Figure 1.**
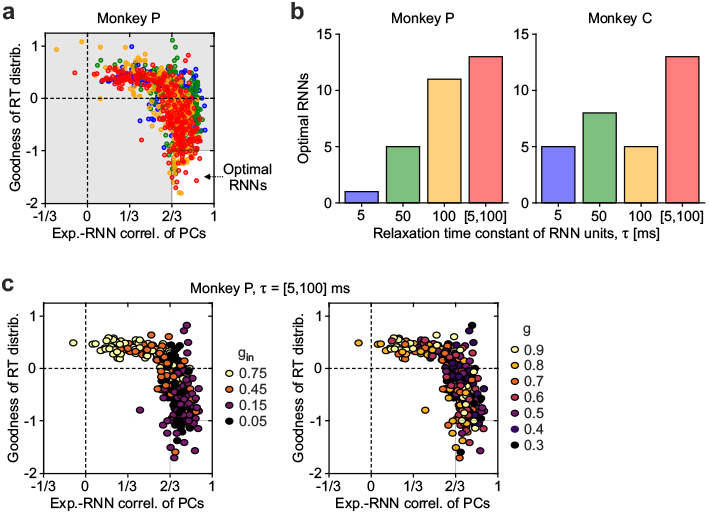
Grid search of optimal RNN hyperparameters. Explored values for the standard deviation of recurrent and input synaptic weights ranged over the intervals *g* ∈ [0.3, 0.9] and *g*_in_ ∈ [0.05, 0.75], respectively. Tested distributions of relaxation time constants *τ* of the RNN units included three homogeneous values (5 ms, 50 ms and 100 ms) and one uniform distribution in the range [5, 100] ms. For each combination of hyperparameters (*τ, g*, and *g*_in_), 10 RNNs were generated with random synaptic matrices and trained to evaluate their performance, resulting in a total of 1,120 networks. **a**, Performance of tested RNNs was assessed based on both the fit of the RT distribution (goodness quantified by the log_10_ of the *p*-value from the Kolmogorov-Smirnov test comparing experimental and digital RTs) and the average correlation between the top three PCs from the experiments and the inferred RNNs. Optimal inferred RNNs were defined as those with *p <* 0.1 and an average correlation coefficient greater than 66%. **b**, Number of optimal RNNs among the tested models for different *τ* distributions in Monkey P (left) and C (right). **c**, Performance of tested RNNs with uniform distribution of *τ* values for Monkey P. Circle colors indicate the standard deviations of the recurrent (*g*) and input (*g*_in_) synaptic weights tested (right and left, respectively).

